# Vcam1+ Fibro-adipogenic Progenitors Mark Fatty Infiltration in Chronic Limb Threatening Ischemia

**DOI:** 10.1101/2024.07.08.602430

**Authors:** Qunsheng Dai, Changxin Wan, Yueyuan Xu, Kaileen Fei, Lindsey A. Olivere, Brianna Garrett, Leo Akers, Derek Peters, James Otto, Christopher D. Kontos, Zhiceng Ji, Yarui Diao, Kevin W. Southerland

## Abstract

Skeletal muscle health and function is a critical determinant of clinical outcomes in patients with peripheral arterial disease (PAD). Herein, we identify fatty infiltration, the ectopic deposition of adipocytes in skeletal muscle, as a histological hallmark of end-stage PAD, also known as chronic limb threatening ischemia (CLTI). Leveraging single cell transcriptome mapping in mouse models of PAD, we identify a pro-adipogenic mesenchymal stromal cell population marked by expression of Vcam1 (termed Vcam1+ FAPs) that expands in the ischemic limb. Mechanistically, we identify Sfrp1 and Nr3c1 as regulators of Vcam1+ FAP adipogenic differentiation. Loss of Sfrp1 and Nr3c1 impair Vcam1+ FAP differentiation into adipocytes *in vitro*. Finally, we show that Vcam1+ FAPs are enriched in human CLTI patients. Collectively, our results identify a pro-adipogenic FAP subpopulation in CLTI patients and provide a potential therapeutic target for muscle regeneration in PAD.

## Introduction

Peripheral arterial disease (PAD), which affects more than 230 million individuals worldwide, represents a significant global healthcare problem^1^. The most severe clinical manifestation of PAD is chronic limb threatening ischemia (CLTI), which presents as rest pain, non-healing ulcers, or gangrene^2^. The primary therapy for CLTI is revascularization^3^. However, despite significant advancements in revascularization techniques, the outcomes for CLTI still remain quite poor, with major amputation rates at 25% at 1 year^4^. These poor outcomes have highlighted the need for a greater understanding of the pathobiology of CLTI and the development of novel therapies.

Recent scientific reports have emphasized the importance of skeletal muscle regeneration as a key determinant of limb outcomes in PAD ^5–7^. In fact, CLTI patients can be biologically distinguished from those with milder forms of PAD by muscle specific signatures ^8–10^. In particular, a radiographic hallmark of the CLTI limb is fatty infiltration, the ectopic deposition of intramuscular adipose tissue at the expense of regenerating muscle fibers ^11–13^. This process leads to progressive muscle dysfunction, impaired ambulation, and ultimately significant quality of life impairments ^14,15^. To date there are no therapeutic strategies to ameliorate fatty infiltration and improve skeletal muscle repair in the ischemic limb. This is in part due to an incomplete characterization of the tissue and cellular processes that contribute to fatty infiltration in the CLTI limb.

The radiographic identification of adipocytes in the CLTI limb implicates a muscle-resident mesenchymal cell population, term fibroadipogenic progenitor (FAPs), as a critical regulator in the pathobiology of CLTI ^16,17^. FAPs are a diverse cell population that display context-dependent behavior. Under healthy conditions that permit regeneration, FAPs provide regenerative cues that orchestrate muscle stem cell (MuSC) activation, proliferation and differentiation ^18–20^. However, in pathologic conditions, FAPs differentiate into either fibroblasts and/or adipocytes, resulting in replacement of muscle with non-contractile tissue and loss of muscle function ^21–23^. Recent studies have identified FAP subpopulations that contribute to fatty infiltration and have provided insights into cellular mechanisms responsible for this process ^24,25^. However, these studies were performed in experimental models with limited clinical applicability. Hence, the mechanisms that underlie the adipogenic differentiation of FAPs in the CLTI limb remain unresolved. Identification of such molecular processes may provide critical insights that would facilitate the development of muscle-specific therapies for CLTI patients.

Accordingly, in this study, we sought to better understand the molecular and cellular mechanisms responsible for adipogenic differentiation of FAPs in the CLTI limb. First, using skeletal muscle specimens from human PAD patients, we validate prior clinical findings, and demonstrate transcriptionally and histologically that fatty infiltration is a signature of the CLTI limb. Next, we used a mouse model that phenocopies CLTI, to identify Vcam1+ FAPs as an injury-induced population with increased adipogenic potential. Furthermore, our studies demonstrate that Sfrp1 and Nr3c1 are required for adipogenic differentiation of Vcam1+ FAPs, and finally we show that FAPs in CLTI patients are enriched in Vcam1, Sfrp1, and Nr3c1. These findings support a potential novel mechanism responsible for fatty infiltration and failed muscle regeneration in CLTI.

## Methods

### Human tissue collection

Paired human skeletal muscle samples were obtained from patients undergoing limb amputation for PAD^31^. Tissue was embedded in OCT and 8-μm sections were prepared for histologic analysis.

### Preclinical PAD model

C57BL/6 and BALB/c mice age 10-12 weeks were anesthetized with isoflurane. Hindlimb ischemia was induced by surgical ligation of the femoral artery as previously described ^31,48^. Laser doppler perfusin imaging (LDPI) was performed with a Moor Instrument LDI2-High Resolution (830nM) System (Moor, Axminster, UK) to quantify perfusion.

### Oil Red O staining on tissue sections (human and mice)

The mouse tibialis anterior skeletal muscle from day 14 hindlimb ischemia was harvested and the human skeletal muscle from both ischemic (distal) and non-ischemic (proximal) parts were obtained from surgical amputation specimens, all the tissues were embedded in OCT compound in liquid N2. Cryostat sections (8 μm) were prepared on microscope slides (Fisher Scientific, Superfrost Plus) for histological analysis.

Frozen sections were allowed to come to room temperature (RT) for 10 minutes then fixed by 10% neutral buffered formalin for 10 minutes at RT followed by incubation in propylene glycol for 5 minutes at RT and in heated (60°C) Oil Red O solution for 20 minutes using Oil Red O Stain Kit (Abcam, ab150678) according to the manufacturer’s protocol. Slides were observed and images were acquired using a Zeiss Axio Imager Z2 Upright Microscope at x 100 magnification at the Duke Light Microscope Core Facility. Lipid with Oil Red O positive area was visualized and measured using the ImageJ (NIH) software.

### Isolation and digestion of hindlimb

Hindlimb ischemia (HLI) was performed as previously described by our group on C57BL/6 mice aged 8-12 weeks. At 3 days post-ischemia, hindlimb muscles were surgically dissected, harvested, dissociated, and digested with Collagenase II (ThermoFisher Scientific, 17101015). Briefly, the isolated hindlimb muscles were minced finely and then subjected to enzymatic digestion for 1.5 hours at 37°C. The muscle dissociation buffer contained collagenase II (700-800 U/mL) and dispase (11 U/mL) in Ham’s F-10 nutrient mixture with 1 mM l-glutamate (ThermoFisher Scientific, 11550043) supplemented with 10% horse inactivated serum (HS) (Gibco) plus penicillin-streptomycin (P/S) (Gibco). Upon dissociation, the cell suspension was subject to filtration through a 40 mm cell strainer (Corning, 352340) and centrifugation for 5 min at 500 g at 4°C.

### FACS isolation of fibro-adipogenic progenitor cells (FAPs)

The single-cell suspension was diluted to 1 x 105 cells/mL with a washing medium (WM: Ham’s F-10 nutrient mixture with 1 mM l-glutamate supplemented with 10% HS plus P/S). Cells were stained with 7-AAD to assess viability and were incubated with the primary antibodies in a rotating shaker at 4°C for 45 minutes protected from light. The following antibodies were used to isolate two subgroups of FAPs: (1) CD31-APC (-), CD45-APC (-), Sca1-FITC (+), VCAM-1 biotin PE-Cy7 (-), and a7-Integrin-PE (-), (2) CD31-APC (-), CD45-APC (-), Sca1-FITC (+), VCAM-1 biotin PE-Cy7 (+), and a7-Integrin-PE (-). Sorting was performed on a BD DIVA sorter at the Duke Flow Cytometry core facility. Data were collected and analysis was performed using FlowJo software. Strict gating schemes were applied to prevent nonspecific cellular contamination and fractionated cells were collected in PBS + 2% FBS + P/S or WM.

### Mouse FAP cell culture and adipogenesis assay

FACS-sorted FAPs were cultured and grown on ECM (Sigma, E1270) coated plates in growth media (GM) consisting of DMEM (ThermoFisher Scientific, 11995065) supplemented with 10% HS, 20% FBS, P/S, and 5 ng/mL bFGF (ThermoFisher Scientific, PHG0367) at 37°C and 5% CO2. GM was changed every three days until about 80% cell confluence. For adipogenic induction, when FAPs growth reached 60-80% confluence, the GM was replaced with StemPro® adipogenesis Differentiation Medium (Gibco, StemPro® adipogenesis Differentiation Kit, A10070-01) and were maintained for 3 and 6 days, respectively. At the endpoint of differentiation, cells were fixed and stained with Oil Red O and antibodies for immunofluorescence staining for evaluating adipogenesis.

### Oil Red O staining in cultured cells

For visualization and quantification of Oil Red O staining, FACS-sorted subgroup FAPs (Vcam1+ and Vcam1-FAPs) were grown in a 12-well ECM-coated plate. The Oil red O solution (Sigma-Aldrich) was used for the detection of neutral lipid droplets in adipocytes differentiated from FAPs. The stock solution (0.3% solution of Oil red O in 100% isopropanol) was dissolved in ddH2O in 3:2 ratios. The cells were fixed in 10% neutral formalin for 60 minutes at room temperature (RT) followed by two washings with ddH2O then stained with Oil Red O solution for 20 minutes at RT, hematoxylin was used for counterstaining. Lipid droplets appeared red and nuclei appeared blue. Images were acquired using an Olympus IX70 microscope at x 200 magnification. After finishing Oil Red O staining and image acquisition, staining was extracted in 300 uL/well 100% isopropanol, and 200 uL was used to measure Oil Red O stain in a 96-well plate reader at 492 nm.

### Immunofluorescence staining of cultured cells

FACS-sorted subgroup FAPs (Vcam1+ and Vcam1-FAPs) were seeded and grown in MatTek glass bottom Microwell dishes (MatTek, 35 mm Dish No. 0 Coverslip 10 mm Glass, P35GCOL-0-10-C), adipogenesis was induced as described previously. At the endpoint of differentiation, cells were fixed in 4% paraformaldehyde for 10 minutes and permeabilized with 0.2% Triton X-100 in PBS for 5 minutes and then in PBS for three 5-minute washes. Blocking solution (10% normal goat serum in PBS and/or AffiniPure Fab Fragment Goat Anti-Mouse IgG (H+L) (Jackson ImmunoResearch laboratories, 115-007-003) was applied for 30 minutes at RT followed by primary antibodies [perilipin (Cell signaling, 9349S), PPARg (Cell signaling, 2435S), myosin heavy chain (Abcam, 37484) and myogenin (Abcam, 1835)] incubation overnight at 4°C. The next day, the cells were washed in PBS for three 5-minute washes and then incubated with goat anti-rabbit IgG (H+L) Cross-Adsorbed Secondary Antibody, Alexa Fluor™ 488 (Invitrogen, 11008) or Goat anti-Mouse IgG (H+L) Cross-Adsorbed Secondary Antibody, Alexa Fluor™ 488 (Invitrogen, 11001) for 1 hour at RT. After washing three times with PBS, the cells were counterstained with DAPI for 1 minute at RT. The cells were mounted with Fluoromount-G™ Mounting Medium (Thermo Fischer Scientific, 00-4958-02). Negative control staining included reactions substituting the primary antibody with normal rabbit IgG or normal mouse IgG. Cells were observed and images were acquired using a Zeiss Axio Imager Z2 Upright Microscope at x 200 magnification at the Duke Light Microscope Core Facility.

### siRNA knockdown assay

FACS-sorted Vcam1+ FAP cells were grown to 60-80% confluence and transfected with 40 nM Accell mouse Sfrp1 siRNA SMARTPool (Dharmacon, E-048941-00-0005), 50 nM ON-TARGET plus mouse Nr3c1 siRNA SMARTPool (Dharmacon, L-045970-01-0005) and 50 nM ON-TARGET plus Non-targeting Control siRNAs (Dharmacon, D-001810-04-05) diluted in Opti-MEM™ I Reduced Serum Medium (ThermoFisher Scientific, 11058021) and transfected with Lipofectamine™ RNAiMAX Transfection Reagent (Invitrogen, 13778030) per the manufacture’s recommendation for 48 to 72 hours.

### Sfrp1 treatment and administration of Way-316606

FACS-sorted Vcam1-FAPs were grown to 60-80% confluence followed by adipogenic differentiation and were treated with 500 ng/mL recombinant mouse sFRP-1 protein (R&D systems, 9019-SF) with or without sFRP-1 inhibitor, 2 uM Way-316606 hydrochloride (Tocris, 4767) or vehicle control (DMSO). At the endpoint of differentiation and treatment, the cells were fixed for Oil Red O staining and immunofluorescence staining to assess adipogenesis.

### RNA isolation and real-time quantitative RT-PCR

Total RNA was extracted from cells using TRIzol reagent (Invitrogen), chloroform extraction, and isopropanol precipitation followed by DNase I (Invitrogen, DNA-free™ DNA Removal Kit, AM1906) treatment according to the manufacturer’s protocol. RNA was stored at –80°C. DNase I-treated total RNA (1 μg) was used for first-strand cDNA synthesis (reverse transcription) using SuperScript™ IV VILO™ Master Mix (Invitrogen, 11756050), quantitative real-time RT-PCR was performed using SYBR™ Green Universal Master Mix (Applied Biosystems, 4309155) to assess gene expression. (mouse Sfrp-1 forward primer: CAATACCACGGAAGCCTCTAAGC, mouse Sfrp1 revise primer: GCAAACTCGCTTGCACAGAGATG, mouse Nr3c1 forward primer: TGGAGAGGACAACCTGACTTCC, mouse reverse primer: ACGGAGGAGAACTCACATCTGG, mouse GAPDH forward primer: CATCACTGCCACCCAGAAGACTG, mouse GAPDH reverse primer: ATGCCAGTGAGCTTCCCGTTCAG). The reaction parameters included an initial 10 minutes of denaturing at 95°C followed by 40 cycles of denaturing for 15 seconds at 95°C, annealing for 60 seconds, and extending for 60 seconds at 60°C. After a final extension, melt curve analysis was performed.

### Western blot analysis

Western blot analyses were used to compare protein levels in the cells with different treatments. Total protein from the cultured cells was isolated by RIPA Lysis and Extraction Buffer (ThermoFisher Scientific, 89900) with Halt™ Protease Inhibitor Cocktail (ThermoFisher Scientific, 87786). Total protein concentration was determined by a detergent compatible DC™ Protein Assay Kit II (Bio-Rad Laboratories, #5000112). For Western blot analysis, 30-50 μg protein was separated on a 10% polyacrylamide gel in Tris/glycine/sodium dodecyl sulfate buffer and transferred to polyvinylidene difluoride (PVDF) membrane in Tris/glycine buffer. The membrane was blocked with 5% non-fat milk for 1 hour at RT and incubated overnight at 4°C with primary antibody [Sfrp1 (Invitrogen, MA5-38193), Nr3c1 (Cell signaling, 12041S), b-catenin (Cell signaling, 8814S), GAPDH (Cell signaling, 2118S), and a-tubulin (Cell signaling, 2144S)]. After the primary antibody, membranes were hybridized with the appropriate anti-rabbit IgG HRP-linked secondary antibody (Cell signaling, 7074S) for 1 hour at RT. The membrane was developed with Pierce™ enhanced chemiluminescence (ECL) Western Blotting Substrate (ThermoFisher Scientific, 32209) per the manufacturer’s recommendation and exposed to ChemiDOC™MP Imaging system (Bio-Rad Laboratories).

## Transcriptomic analysis

### Human scRNA-seq analysis

Transcriptome profiles of CLTI patients were obtained from the Gene Expression Omnibus (GEO;GSE227077) from Southerland et al ^31^.

### Mouse scRNA-seq analysis

The sequencing reads (10× Genomics) were processed using the Cell Ranger pipeline (v3.1.0,) with GRCm38 as the reference genome. Downstream analysis was performed using R package Seurat (v4.3.0) with the output read count matrix as input^49^. We filtered out cells using feature number > 4100 or < 1000, or mitochondrial RNA ratio > 25%. Doublets were identified and removed using Python package Scrublet (v0.2.3)^50^. Raw read count matrix was normalized and log-transformed by the NormalizeData function. Top 2000 highly variable features were selected using FindVariableFeatures function and then scaled by ScaleData function. Principle component analysis was performed using RunPCA function with top 50 pcs saved. For multi-sample integration, the harmony algorithm was performed using the RunHarmony function with PCA result as input ^51^ Using harmony embeddings, we then compute UMAP and cluster cells. Cell type annotations of each cluster were made based on these marker genes. FAPs were selected after annotation and processed from read count matrix using the steps mentioned above. Marker genes of each cluster were computed using the function FindAllMarkers.

### Mouse bulk RNA-seq analysis

Bulk RNA-seq reads were aligned to the GRCm38 reference genome by STAR^52^. Read count matrix for each sample was computed using FeatureCounts (v2.0.1)^53^. The DEGs were identified using R package DESeq2 (v1.34.0) with a threshold of adjusted p-value less than 0.05 and fold change greater than 2^54^. GO enrichment analysis was performed with these DEGs via R package clusterProfiler (v4.8.3)^55^.

General statistics

## Results

### Human CLTI limb has an adipogenic phenotype

Computed tomography (CT) analyses have established fatty infiltration as a feature of the CLTI limb^11,12,26^. We sought to biologically validate these clinical findings. First, we used a publicly available human PAD skeletal muscle bulk RNA-seq dataset, which contains samples from non-PAD (n=15), intermittent claudication (mild PAD) (n=20) and CLTI (n=16) patients to assess the gene expression of adipocyte marker genes^9^. We found a clear upregulation of adipocyte marker genes in CLTI patients compared to intermittent claudication and non-PAD patients (Figure 1a-d). Next, we obtained paired proximal and distal muscle specimens from 8 CLTI patients undergoing limb amputation (Table S1). In this experimental design, proximal specimens are from non-ischemic muscle while the distal specimens are ischemic (Figure 1e). Furthermore, this approach controls for heterogeneous clinical variables, such as age, diabetes, and medication use that can influence adipogenesis. To assess adipogenesis, we used the neutral lipid stain Oil Red O (ORO). Proximal (non-ischemic) muscle specimens displayed minimal ORO staining. In contrast, extensive ORO staining was noted in distal (ischemic) muscle specimens (Figure 1f, g). These transcriptomic and histologic data indicate that CLTI patients have an adipogenic phenotype.

**Figure 1:**
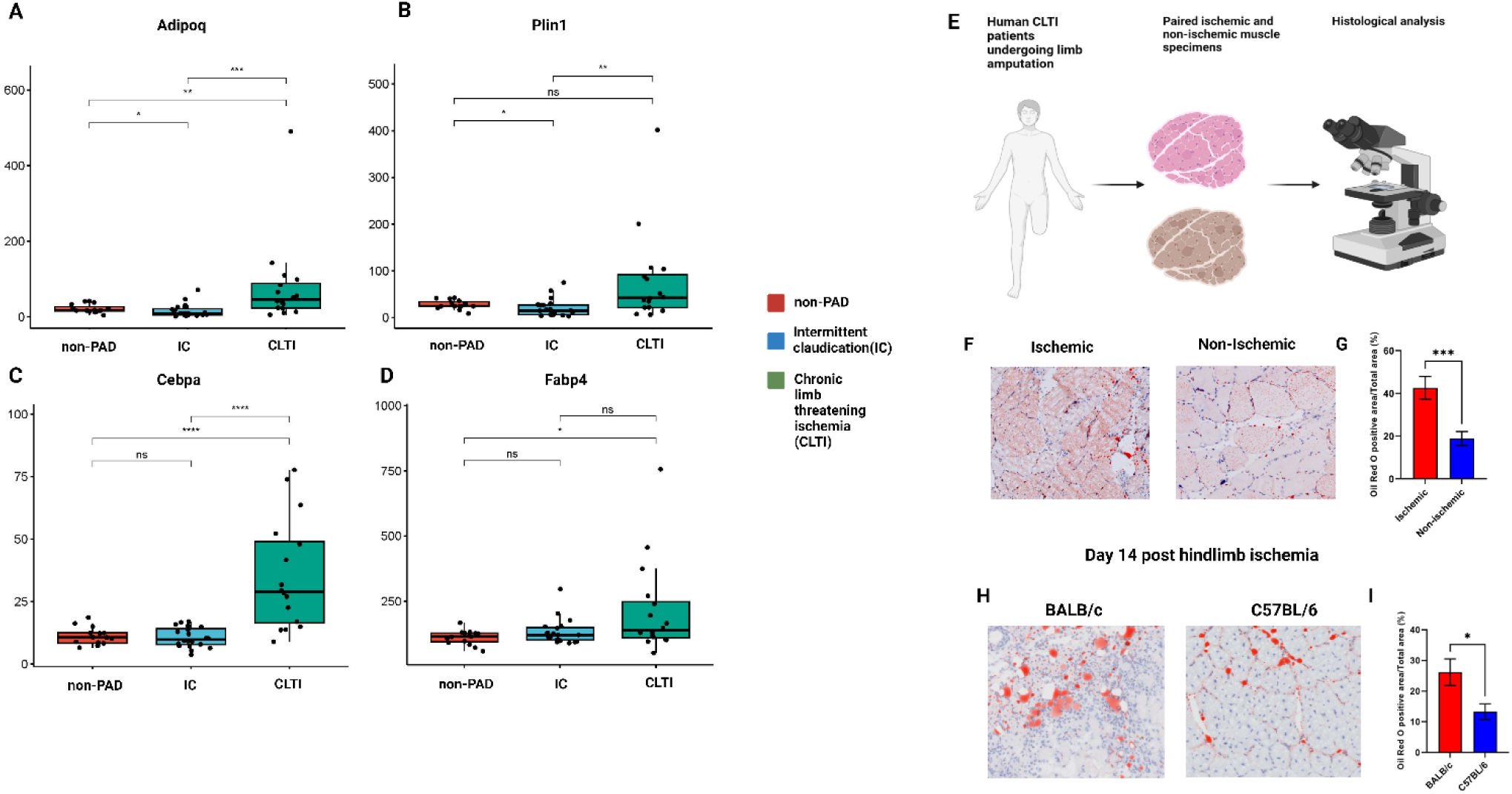
Human CLTI limb has an adipogenic phenotype. A-D: mRNA expression of Adipoq, Plin1, Cebpa, Fabp4 determined by human PAD bulk RNA-seq data set (n= 51). A one-way ANOVA with Holm-Sidak’s multiple comparison test (*p<0.05, **p<0.01, ***p<0.001) was used. E: Experimental design for human histologic analysis. Paired proximal and distal muscle specimens form human CLTI patients (n=8) F: Representative images of immunostaining of ORO in ischemia and non-ischemic muscle specimens in human CLTI patients (n=7) G: Quantification of ORO staining. Data shown as mean ± SEM, ***p<0.001 H: Representative images of immunostaining of ORO in ischemic muscle on day 14 status-post HLI in BALB/c (n=7) and C57BL/6 (n=9) mice I: Quantification of ORO staining. Data shown as mean ± SEM, *p<0.05

We then sought to identify a suitable mouse model to elucidate the mechanisms of adipogenesis in the ischemic limb. To this end, we performed femoral artery ligation to induce hindlimb ischemia (HLI), a model of PAD, on both BALB/c and C57BL/6 mice, two commonly used models to study ischemic responses. Tibialis anterior (TA) muscle from the ischemic limb was obtained at day 14 post HLI and stained for ORO. Compared to that from C57BL/6 mice, which have robust regenerative capacity ^27–29^, muscle collected from BALB/c mice demonstrated significantly more adipogenic replacement, resembling that in the ischemic muscle from human patients with CLTI (Figure 1h, i). These data, together with previous reports showing that BALB/c mice recapitulate many key aspects of the human CLTI phenotype, such as impaired muscle regeneration and ischemic tissue loss ^28–31^, support the use of this model model to investigate mechanisms of the pathologic adipogenesis in the ischemic limb.

### Single-cell RNA sequencing reveals Vcam1+ FAPs as a candidate adipogenic population in the ischemic limb

To explore the cellular mechanisms of fatty infiltration in an unbiased, high-resolution manner, we generated a dynamic single cell atlas of successful muscle regenerative (C57BL/6 mice) and impaired muscle regeneration (BALB/c mice). We collected ischemic TA muscles of 10-12 week old mice at days 0 (no injury), 1, 3, 7, 10, and 14 after HLI (n = 2-3 mice per time point) (Figure 2a). We enzymatically digested TA muscles into single-cell suspensions and performed scRNA-seq on the 10x Chromium platform. We captured 159,189 cells and identified 15 unique clusters (Figure S1a-d). Next, we re-clustered all the FAPs (n = 37,260) in this dataset on a new UMAP space. We identified 12 distinct FAP clusters in both mouse strains and across all time points (Figure 2b). To identify a cluster responsible for the adipogenic phenotype, we first turned our attention to day 14, the time point at which BALB/c mice displayed significantly more ORO staining (i.e. adipogenesis) than C57BL/6 mice (Figure 1d,e). On day 14 in BALB/c mice, there were three predominant FAP sub-clusters, clusters 0 (24.5%),1 (23.5%), and 2 (23.5%).

**Figure 2:**
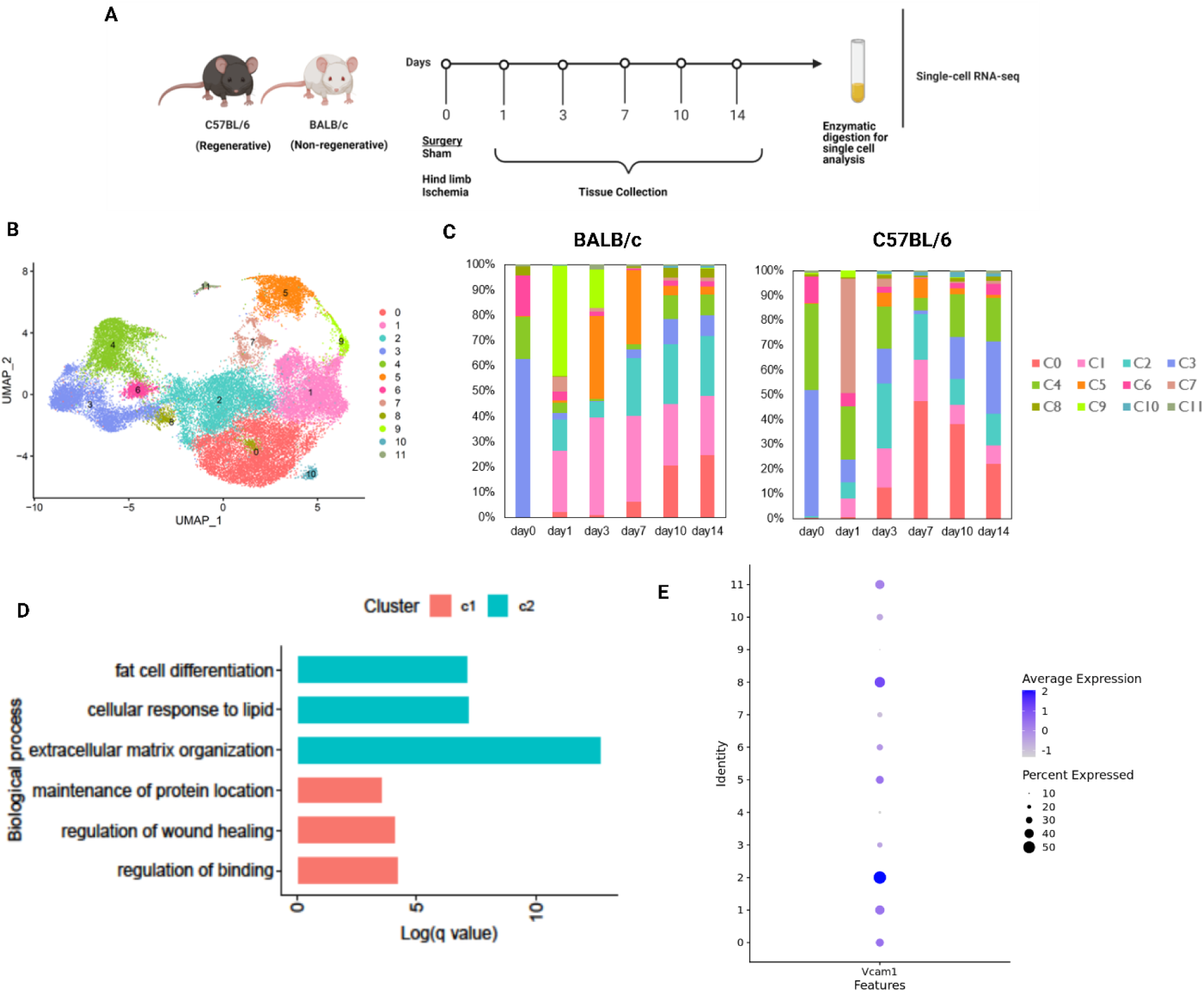
scRNA-seq in a mouse PAD model reveals Vcam1+ FAPs as a candidate adipogenic population. A: Experimental design for generation of scRNA-seq atlas (n = 2 biological replicates per strain at each time point). B: Uniform Manifold Approximation Projection (UMAP) visualization of FAP clusters colored by identify (n = 37,260) C: Percentage of FAP populations by mouse strain and time point D: Bar plots showing enriched pathways from GO terms of FAP cluster 1 vs cluster 2 E: Dot plot demonstrating increased Vcam1 expression in FAP cluster 2 compared to all other FAP clusters (adjusted p-value is 4.133265e^-236^ and log2 fold change is 0.5454154).

In contrast, clusters 1 (C1) and 2 (C2) represented a small fraction, 7.2 and 12.9% respectively, of the day 14 FAP population in C57BL/6 mice (Figure 2c); hence, we reasoned that C1 and C2 were reasonable candidates driving adipogenesis. Pathway analysis of C1 and C2 gene profile revealed the enrichment of genes involved in fat cell differentiation in C2 compared to C1 (Figure 2d). Thus, C2 appeared to be the most suitable adipogenic candidate cluster. Next, we sought to identify a cell-surface marker for this cluster that would permit FACS isolation of these cells for *in vitro* characterization of their functional properties. Notably, Vcam1 has been shown to be a marker of injured FAPs^32^. Accordingly, in our data we found that Vcam1 expression was upregulated in C2 compared to other FAP subclusters, supporting its use as a marker gene for this population (Figure 2e).

### Vcam1+ FAPs display increased adipogenic potential and lipid metabolism

To better characterize Vcam1+ FAPs and gain mechanistic insights into their role in adipogenesis, we generated a bulk RNA-seq dataset of Vcam1+ and Vcam1-FAPs isolated from the ischemic limb of C57BL/6 mice and cultured in adipogenic differentiation media conditions for 3 days (Figure 3a; Figure S2a). We chose this time point to capture the transcriptional changes preceding adipogenic differentiation since both Vcam1+ and Vcam1-FAPs display minimal adipogenic differentiation at this time point (Figure S2b-g). Principal component analysis demonstrated that Vcam1+ and Vcam1-FAPs clustered separately (Figure 3b). Compared to Vcam1-FAPs, Vcam1+ FAOs expressed 587 upregulated and 575 down-regulated genes (Figure 3c), which are highly enriched for fatty acid and lipid metabolism (Figure 3d).

**Figure 3:**
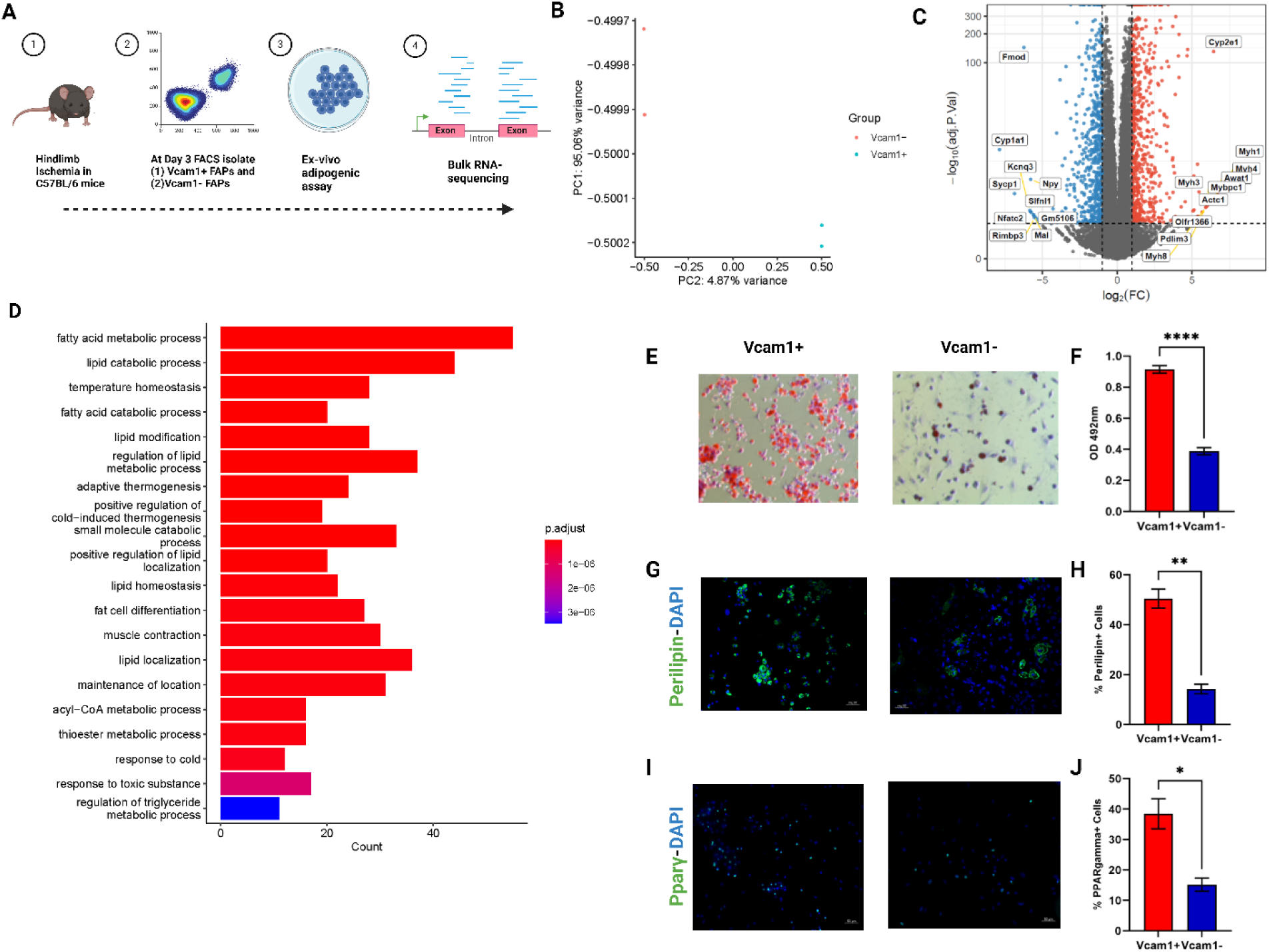
Vcam1+ FAPs display increased adipogenic potential and lipid metabolism. A: Experimental design for bulk RNA-seq dataset. 2 biologic replicates B: Principal component analysis (PCA) showing sample distances from mouse bulk RNA-seq data comparing day 3 Vcam1+ and Vcam1-FAPs, color-coded by FAP subpopulation C: Volcano plot demonstrating significantly upregulated (red) and downregulated (blue) genes in Vcam1+ versus Vcam1-FAPs in adipogenic media for 3 days (log2FC≥1 and p<0.05) D: Dot plot showing upregulated pathways in Vcam1+ versus Vcam1-FAPs E: Representative images of immunostaining for ORO in adipogenic media for 6 days F: Quantification of ORO staining. Data shown as mean ± SEM, ***p<0.001 G: Representative images of perilipin (green) and DAPI (blue) staining H: Quantification of perilipin staining. Data shown as mean ± SEM, **p<0.01 I: Representative images of Pparγ (green) and DAPI (blue) staining J: Quantification of Pparγ staining. Data shown as mean ± SEM, *p<0.05

We further investigated the adipogenic capacity of Vcam1+ FAPs. After culturing in adipogenic conditions for 6 days, Vcam1+ FAPs displayed significantly greater adipogenesis compared to Vcam1-FAPs, evident by increased ORO, perilipin, and PPARγ staining (Figure 3e-j). Taken together, these data demonstrate that Vcam1+ FAPs are pro-adipogenic subcluster.

### Sfrp1 regulates Vcam1+ FAP adipogenic differentiation

To probe for the underlying mechanism driving Vcam1+ FAP-promoted adipogenesis, we explored the RNA-seq profiles of Vcam1+ and Vcam1-FAPs for Wnt signaling, which has been reported to be a critical inhibitor of adipogenesis^33,34^. Indeed, Wnt signaling pathways were significantly downregulated in Vcam1+ FAPs compared to Vcam1-FAPs (Figure 4a). Consistently, Sfrp1, an extracellular ligand that inhibits canonical Wnt signaling, was enriched in Vcam1+ FAPs compared to Vcam1-FAPs in adipogenic media (Figure 4b). The positive association between Sfrp1 and Vcam1 is also supported by the co-expression patterns of these two genes in scRNA-seq data, with a Pearson correlation of 0.52 (Figure 4c). Furthermore, Sfrp1 was enriched in cluster 2, the Vcam1 high cluster (Figure S3a). While Sfrp1 expression was higher in Vcam1+ FAPs, both at the mRNA and protein levels, the expression of B-catenin, a marker of canonical Wnt signaling, was lower (Figure S3b, Figure 4d). Collectively, these data support a strong causal relationship between Sfrp1, inhibition of Wnt signaling, and adipogenesis in Vcam1+ FAPs.

**Figure 4:**
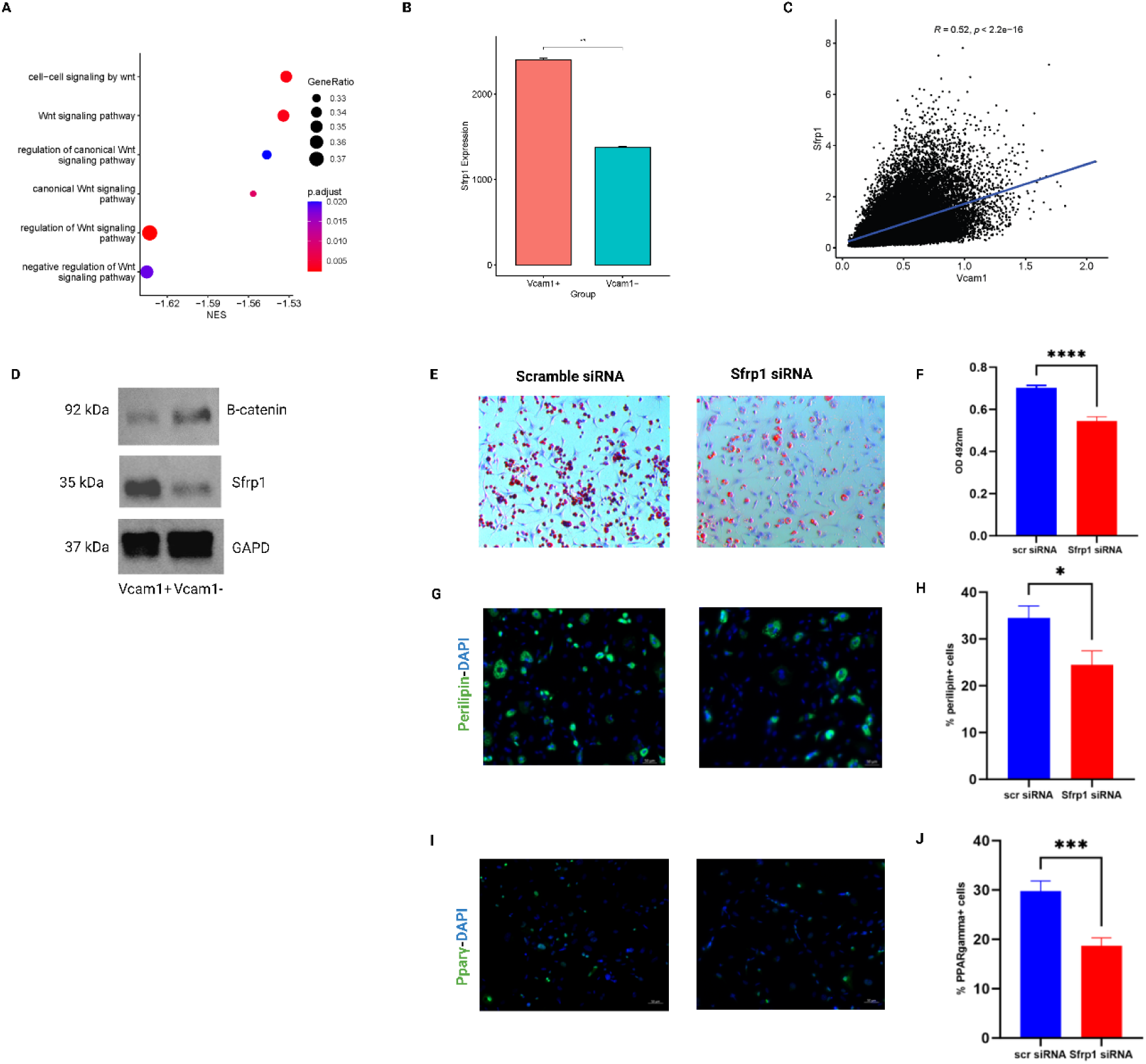
Sfrp1 regulates Vcam1+ FAP adipogenic differentiation. A: Dot plot demonstrating Wnt signaling pathways in Vcam1+ versus Vcam1-FAPs B: Bar graph demonstrating Sfrp1 expression in Vcam1+ versus Vcam1-FAPs C: Heatmap representing Pearson’s correlation values of Sfrp1 and Vcam1 expression in FAPs D. β-catenin and Sfrp1 protein expression in cell lysates from Vcam1+ and Vcam1-FAPs. GAPDH was used for a loading control and signal normalization (n = 3 different experiments) E: Representative images of ORO staining of Vcam1+ FAPs treated with siRNA against Sfrp1 and a control siRNA F: Quantification of ORO staining. Data shown as mean ± SEM, ***p<0.001 G: Representative images of perilipin (green) and DAPI (blue) staining of Vcam1+ FAPs treated with siRNA against Sfrp1 and a control siRNA H: Quantification of perilipin staining. Data shown as mean ± SEM, *p<0.05 I: Representative images of Pparγ (green) and DAPI (blue) staining of Vcam1+ FAPs treated with siRNA against Sfrp1 and a control siRNA J: Quantification of Pparγ staining. Data shown as mean ± SEM, ***p<0.001

Next, we determined whether Sfrp1 was sufficient and/or necessary for adipogenic differentiation of FAPs. We FACS-isolated Vcam1+ FAPs and transfected them with a small interfering (siRNA) to knockdown Sfrp1. As a result of effective Sfrp1 knockdown (over 80% at the mRNA level (Figure S3c)), the Vcam1+ FAPs exhibited a significant decrease in adipogenic capacity (Figure 4e-j). In a complementary gain-of-function experiment, we treated the FACS-isolated Vcam1-FAPs (low adipogenic cell population) with recombinant Sfrp1. Compared to vehicle control, exogenous Sfrp1 increased adipogenesis of Vcam1-FAPs, which was abolished by Way-316606, a small molecular inhibitor of Sfrp1 (Figure S3d-i). These data demonstrate that Sfrp1 helps regulate Vcam1+ FAP adipogenic differentiation.

### scRNA-seq and scATAC-seq identify Nr3c1 as a candidate cell-type specific transcription factors that regulate the adipogenic transcriptional program in FAPs

To better understand the mechanisms regulating Sfrp1-mediated adipogenesis, we sought to identify putative transcription factors (TFs) responsible for the adipogenic transcriptional program in FAPs. To accomplish this, we employed several computational methods. First, we revisited our scRNA-seq data set and performed a differential gene expression analysis between Vcam1+ and Vcam1-FAPs. Next, we imputed our gene list into Lisa, a computational platform that uses public chromatin accessibility and ChIP-seq data to infer transcriptional regulators ^35^. Through these analyses, we identified several candidate TFs, including Nr3c1, Cebpb, and Med1 that regulate Vcam1+ FAPs (Figure 5a).

**Figure 5:**
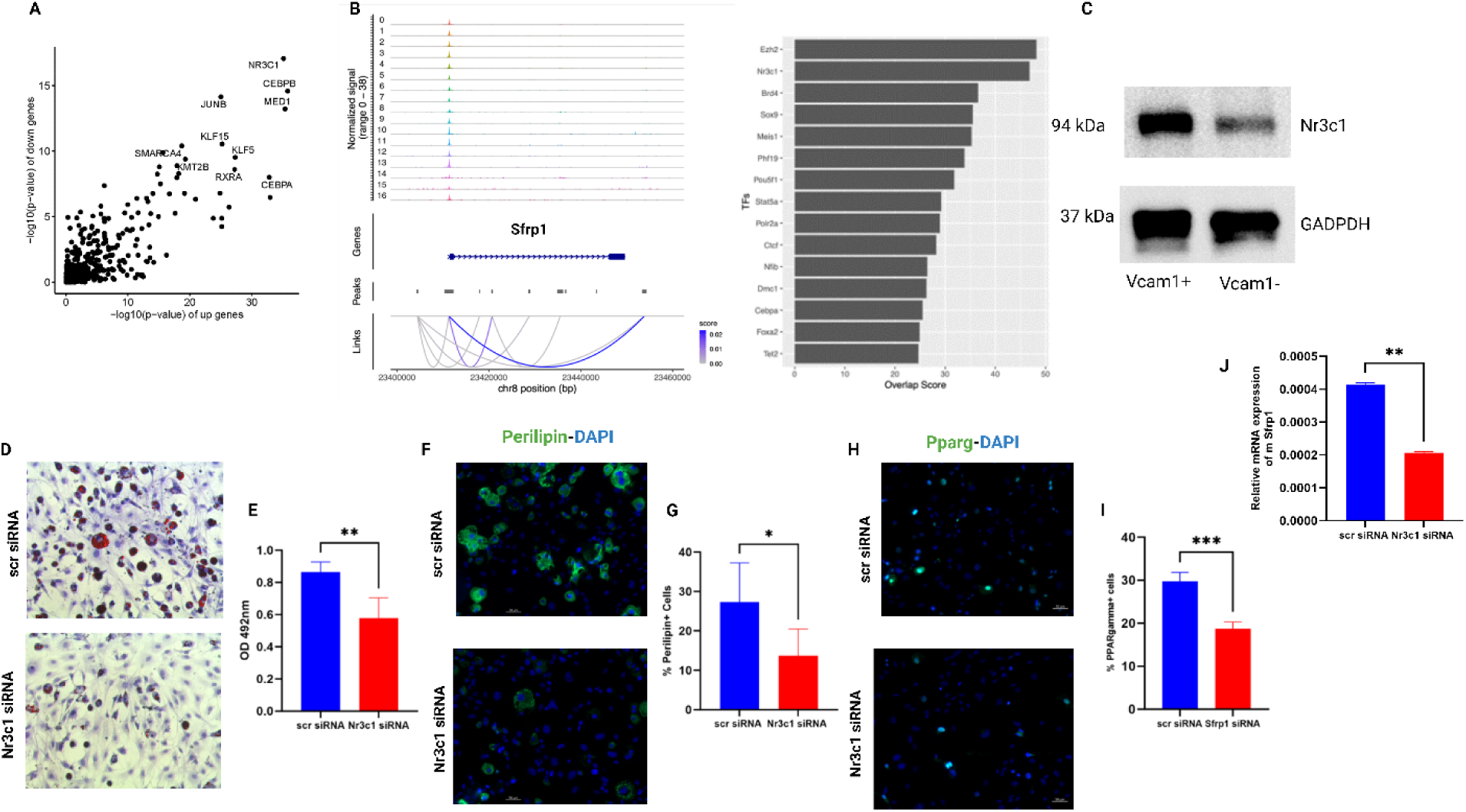
scRNA-seq and scATAC-seq identifies Nr3c1 as a transcription factor that regulates FAP adipogenesis. A: Inferred transcription factors that regulate differential genes of Vcam1+ versus Vcam1-FAPs B: Enhancers with regulation potential to Sfrp1 (left) and transcription factor binding analysis to the enhancers C: Nr3c1 protein expression in Vcam1+ and Vcam1-FAPs in adipogenic media for 3 days D: Representative images of ORO staining in Nr3c1-silenced Vcam1+ FAPs E: Quantification of ORO staining. Data shown as mean ± SEM, **p<0.01 F: Representative images of perilipin (green) and DAPI (blue) staining in Nr3c1-silenced Vcam1+ FAPs G: Quantification of perilipin staining. Data shown as mean ± SEM, *p<0.05 H: Representative images of Pparγ (green) and DAPI (blue) staining in Nr3c1-silenced Vcam1+ FAPs I: Quantification of Pparγ staining. Data shown as mean ± SEM, ***p<0.001 J: Gene expression of Sfrp1 in FAPs treated with siRNA against Nr3c1 and a control siRNA. Student’s t-test (two-tailed, unpaired, **p<0.01)

To further elucidate the regulatory mechanisms underpinning adipogenic differentiation in FAPs, we generated a dynamic chromatin accessibility atlas of regenerative (C57BL/6) and non-regenerative (BALB/c) responses in the ischemic limb at single cell resolution (Figure S4a,b). For the purposes of this analysis, we re-organized all of the FAPs on a new UMAP space. Unbiased clustering revealed 12 distinct FAP sub-populations (Figure S4c). Next, we identified enriched peaks within 10 kilobases of the Sfrp1 locus (Figure 5b) and performed a TF binding analysis, using Cistrome DB Toolkit, to identify TFs with significant binding on these peaks. This analysis also revealed Nr3c1 as one of the candidate TFs for Sfrp1 (Figure 5b). Indeed, Nr3c1 expression was higher in Vcam1+ FAPs compared to Vcam1-FAPs (Figure 5c). Importantly, siRNA knockdown of Nr3c1 (Figure S4d) resulted in a significant decrease in adipogenic differentiation of Vcam1+ FAPs (Figure 5d-i) and Sfrp1 expression (Figure 5j), indicating a role of Nr3c1 in Sfrp1 mediated adipogenic differentiation of Vcam1+ FAPs.

### Human CLTI FAPs share a transcriptional signature with the adipogenic FAP cluster in BALB/c mice

To assess the clinical relevance of our findings, we asked whether FAPs from human CLTI patients displayed a similar gene expression profile as in the adipogenic cluster 2 in BALB/c mice. To do this we used a human CLTI scRNA-seq dataset that we generated previously ^31^. This dataset contains matched proximal/non-ischemic and distal/ischemic skeletal muscle specimens from 3 CLTI patients following limb amputation. After isolating all of the FAPs onto a new UMAP space, we identified 9 distinct FAP clusters, revealing immense transcriptional heterogeneity of FAPs in the human ischemic limb (Figure 6a). Next, we segregated the FAPs by anatomic location in the amputated limb; this revealed disparate transcriptional programs between distal/ischemic and proximal/non-ischemic FAPs (Figure 6b). We then assessed the expression of Vcam1, Sfrp1, and Nr3c1 in these human CLTI FAPs. Vcam1, Sfrp1, and Nr3c1 were all enriched in distal/ischemic FAPs compared to proximal/non-ischemic FAPs (Figure 6c-e). Altogether, these data suggest similar transcriptional signatures characterized by an induction of Nr3c1-Sfrp1 axis in Vcam1+ FAPs in response to muscle ischemia in BALB/c mice and human CLTI patients.

**Figure 6:**
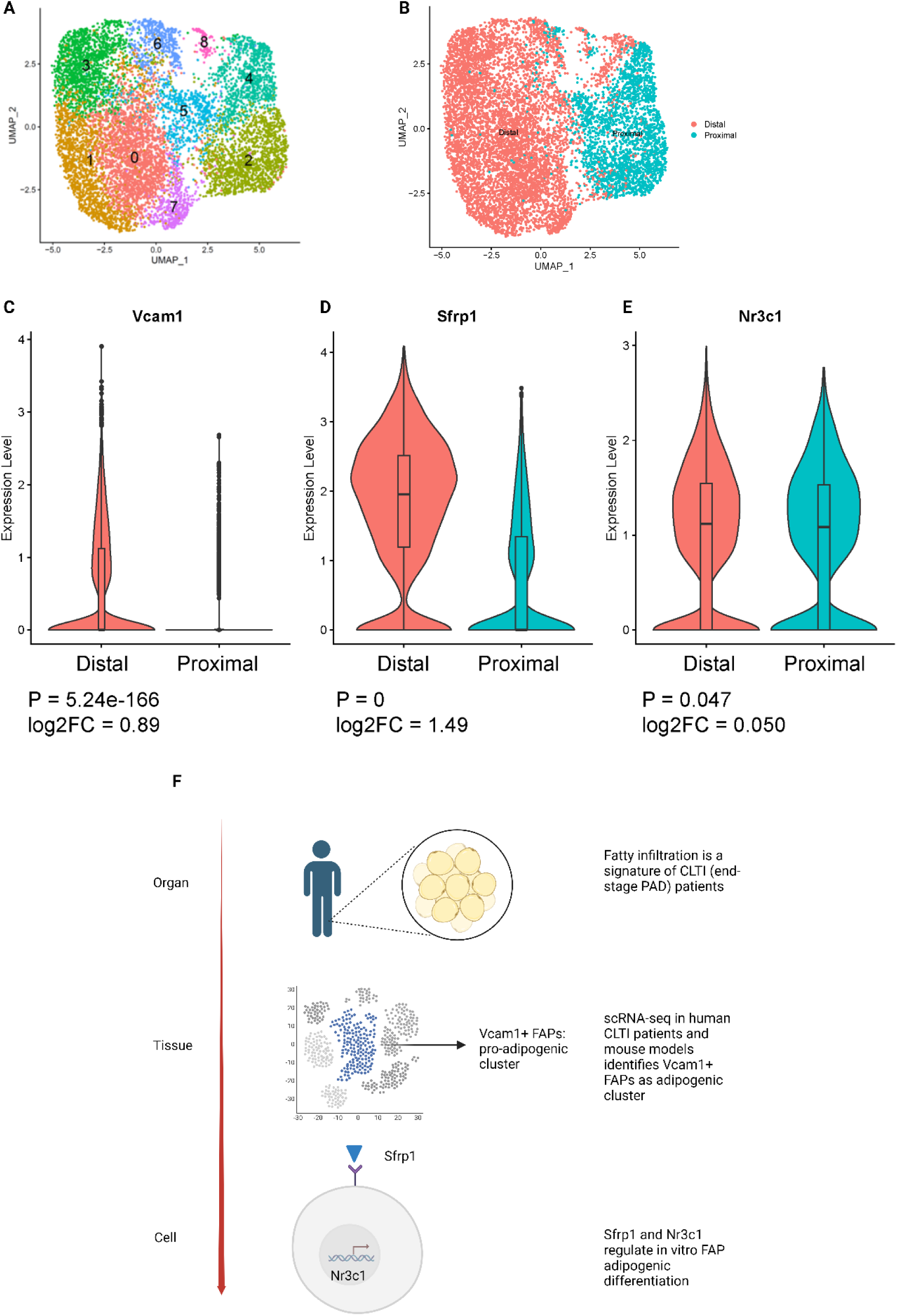
Human CLTI FAPs share a transcriptional signature with BALB/c FAPs. A: UMAP of FAPs in human PAD dataset, color represents subcluster B: UMAP of FAPs in human PAD dataset, color represents location in limb C: Violin plot of Vcam1 expression in distal/ischemic versus proximal/non-ischemic FAPs D: Violin plot of Sfrp1 expression in distal/ischemic versus proximal/non-ischemic FAPs E: Violin plot of Nr3c1 expression in distal/ischemic versus proximal/non-ischemic FAPs F: Schematic summarizing study findings

## Discussion

CLTI represents a grave threat to human life and limb; unfortunately there are a paucity of regenerative therapies for this devastating pathology. Herein, we establish fatty infiltration as a signature of the CLTI limb and identify a pro-adipogenic FAP subcluster. Specifically, (1) we use transcriptional profiling in a murine CLTI model to identify Vcam1+ FAPs as an enriched population in the ischemic limb, (2) we demonstrate that Vcam1+ FAPs display increased adipogenic capacity in part via Sfrp1 mediated inhibition of canonical Wnt signaling; (3) we also identify Nr3c1 as a transcriptional regulator of Vcam1+ FAP adipogenic differentiation; (4) finally, we demonstrate that Vcam1+ FAPs are enriched in the ischemic muscle of CLTI patients (Figure 6f).

Skeletal muscle responses are a key determinant of clinical outcomes in PAD patients^30,36,37^. In fact, CLTI, end-stage PAD, is distinguished from more benign forms of PAD by muscle specific features^8,9,38,39^. In particular, CLTI is hallmarked by the replacement of functional muscle fibers with adipocytes ^11,12^. Ferreira et al. performed an observational prospective study of 116 PAD patients and demonstrated via computed tomography (CT) scans that CLTI patients have more intramuscular fat than intermittent claudication patients and the muscle signature of fatty infiltration was associated with worse clinical outcomes^12^. Sugai et al. obtained CT scans on 327 consecutive patients undergoing revascularization procedures and found that intramuscular fat was associated with major adverse limb events, such as amputation^11^. Our work validates these clinical findings, and our data solidify fatty infiltration as a pathological signature of the CLTI limb, which points to FAPs as a critical element of CLTI pathobiology.

The mechanisms governing FAP-to-adipocyte conversion in the ischemic limb remain unknown. Herein, we uncover a candidate molecular signaling axis that regulates this pathologic cell fate. We identify Vcam1+ as an injury-specific FAP sub-cluster with increased adipogenic potential. This finding is in agreement with prior studies demonstrating Vcam1 as a marker for injury FAPs^32^. We also observed decreased Wnt signaling in Vcam1+ FAPs. Canonical Wnt signaling is a well established inhibitor of adipogenesis ^34^. In a murine muscular dystrophy model, Reggio et al demonstrated that the canonical Wnt/Gsk/B-catenin signaling axis inhibited FAP adipogenic differentiation^40^. Our findings support these conclusions and suggest that Wnt signaling is a conserved mechanism that constrains adipogenesis in PAD as well as other myopathies. Furthermore, we used *in silico* prediction methods along with in vitro inhibition studies to identify Nr3c1 as a candidate upstream transcriptional regulator of the Sfrp1/B-catenin axis. Nr3c1, a nuclear glucocorticoid receptor, is known to promote FAP differentiation in adipocytes. Furthermore, glucocorticoid administration has been demonstrated to augment Sfrp1 expression^41^. Altogether, these findings identify Nr3c1-Sfrp1-B-catenin as a putative molecular axis governing FAP-to-adipocyte conversion, at least in vitro ^42^. Further studies are necessary to validate this mechanism in vivo. However, this pathway represents a potentially targetable mechanism to ameliorate fatty infiltration in the ischemic limb.

The novel finding that Vcam1+ FAPs represent a pathologic FAP subpopulation that promotes adipogenesis in not only a murine model of limb ischemia and human CLTI patients is promising. To our knowledge, this is the first identification of a distinct FAP subpopulation in human CLTI patients. Although several investigators have profiled the transcriptomes of human FAPs at single cell resolution these data sets were derived primarily from healthy skeletal muscle.^7,25,43–45^. An important exception is the work of Farup et al, who identified Thy1+ FAPs as a fibrotic FAP population enriched in type 2 diabetic patients. In addition, Fitzgerald et al used skeletal muscle specimens from gluteus mimimus and rectus femoris muscle in patients with symptomatic hip osteoarthritis undergoing total hip arthroplasty and identified Mme+ FAPs as an adipogenic FAP subpopulation in this context ^25^. Interestingly, in our data set, Mme expression was enriched in the proximal/non-ischemic tissue compared to the distal/ischemic skeletal muscle. These data suggest that disease-specific niches differentially affect FAP heterogeneity, and they highlight the critical importance of disease-specific datasets for biomedical discovery.

In summary, we have identified a pro-adipogenic FAP population defined by Vcam1 expression in the ischemic limb. We demonstrated that the adipogenic differentiation of Vcam1+ FAPs is, in part, regulated by Nr3c1 and Sfrp1. Moreover, we found that this population is enriched in human CLTI patients. Targeting Vcam1+ FAPs via inhibition of the Nr3c1-Sfrp1 axis may represent a novel strategy to mitigate maladaptive adipogenesis in the ischemic limb of CLTI patients.

## Limitations

It is important to note some limitations of our study. First, we have demonstrated the increased ex-vivo adipogenic capacity of Vcam1+ FAPs. These studies do not account for the influence of the skeletal muscle niche on FAP cell fate, which is known to play a critical role in FAP phenotypes ^46,47^. Second, it is quite promising that our preclinical data correlated with fatty infiltration and Vcam1 expression in CLTI patients. However, to demonstrate the ultimate clinical significance of this pathway will require additional in vivo mechanistic studies.

## Supplemental Figures and Tables

**Supplementary Figure 1:**
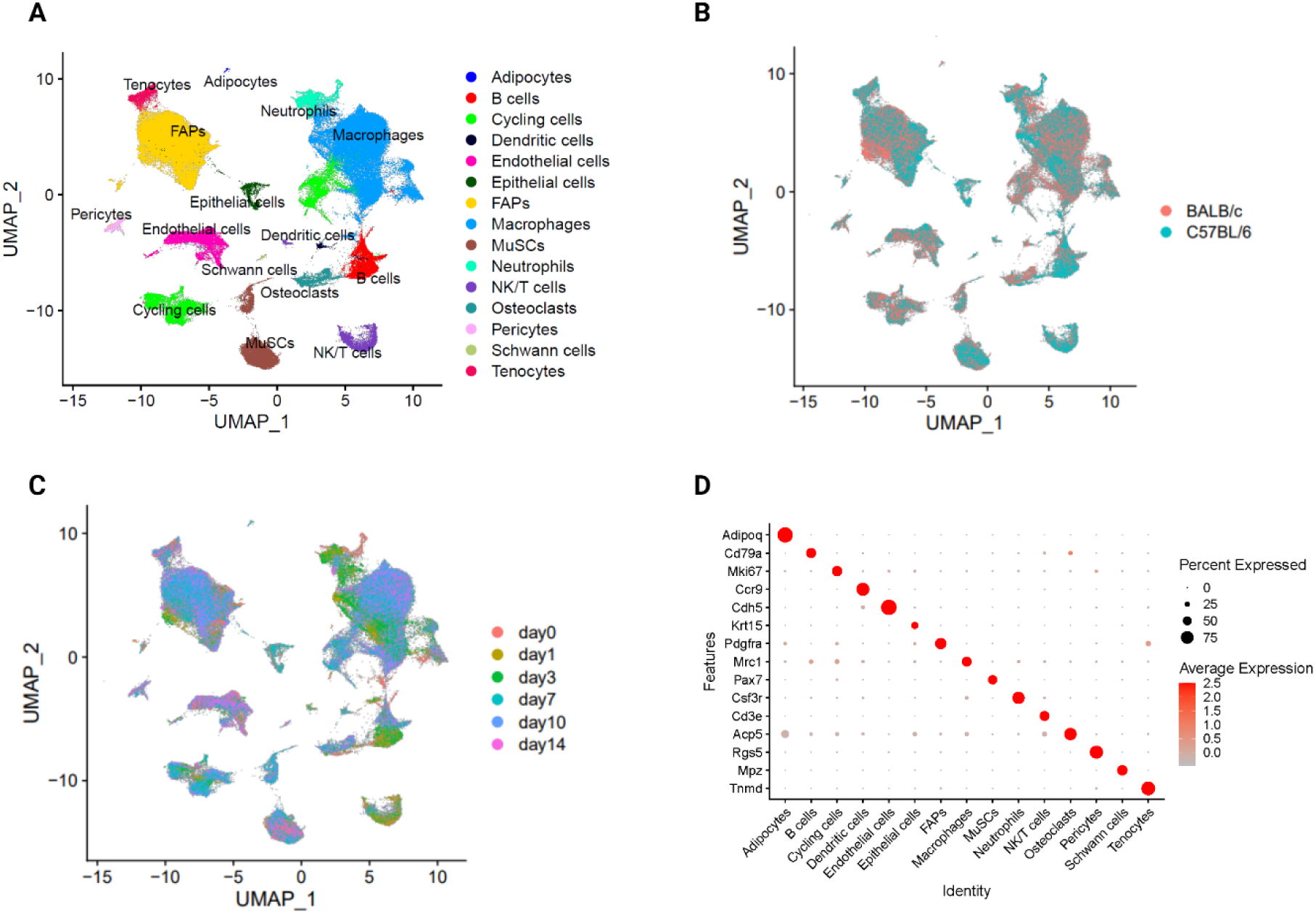
Mouse hindlimb ischemia scRNA-seq atlas (Related to Figure 2) A: UMAP visualization of all cells, colored by cell type B: UMAP visualization of all cells, colored by mouse strain C: UMAP visualization of all cells, colored by time point D: Dot plot demonstrating cell-type marker gene expression

**Supplementary Figure 2:**
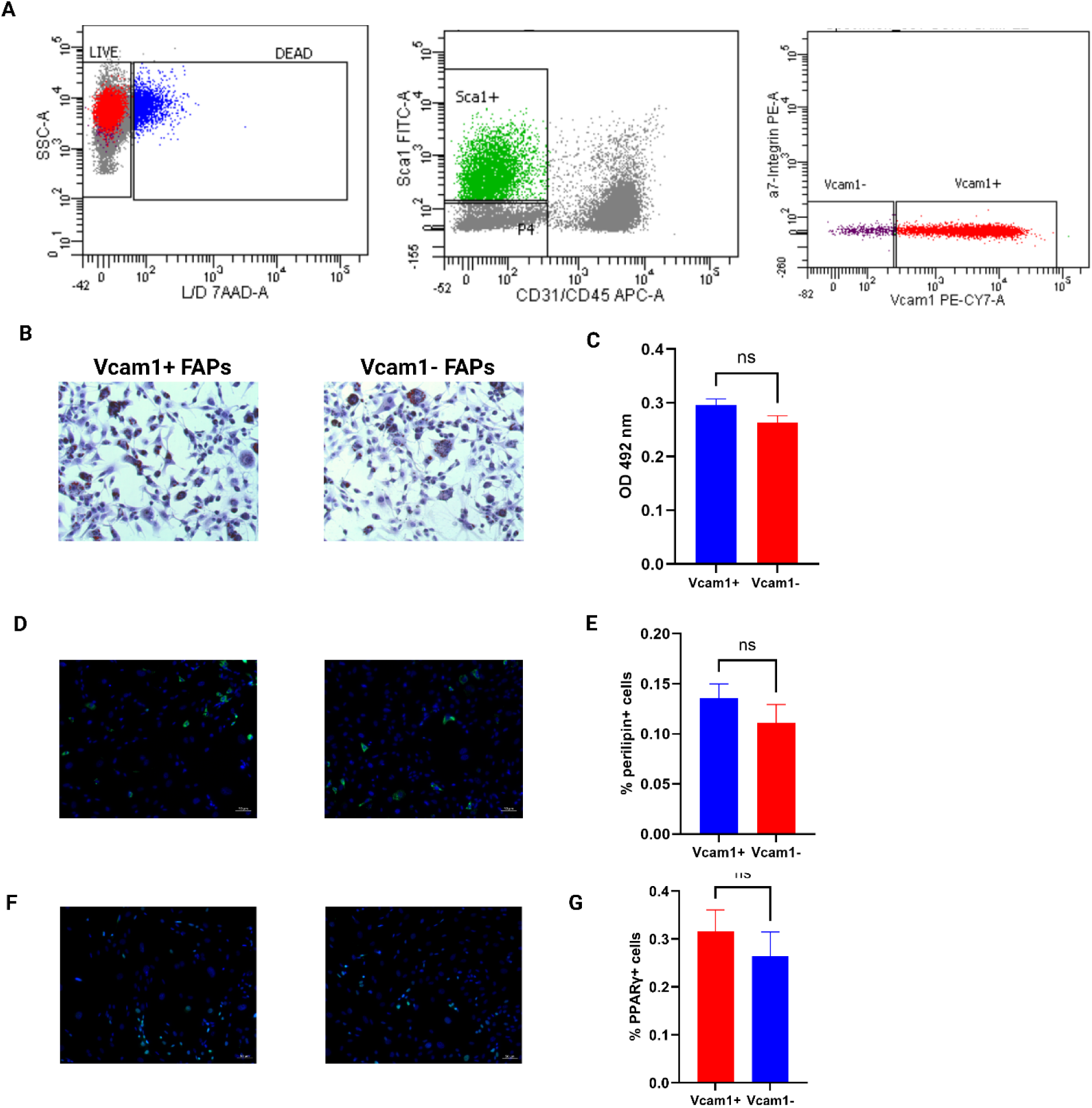
In vitro adipogenic differentiation of Vcam1+ versus Vcam1-FAPs at early time point (Related to Figure 3) A: Representative FACS plot of Vcam1+ and Vcam1-FAPs B: Representative images of immunostaining for ORO in adipogenic media for 3 days C: Quantification of ORO staining. Data shown as mean ± SEM, ns > 0.05 D: Representative images of perilipin (green) and DAPI (blue) staining E: Quantification of perilipin staining. Data shown as mean ± SEM, ns > 0.05 F: Representative images of Pparγ (green) and DAPI (blue) staining G: Quantification of Pparγ staining. Data shown as mean ± SEM, ns > 0.05

**Supplementary Figure 3:**
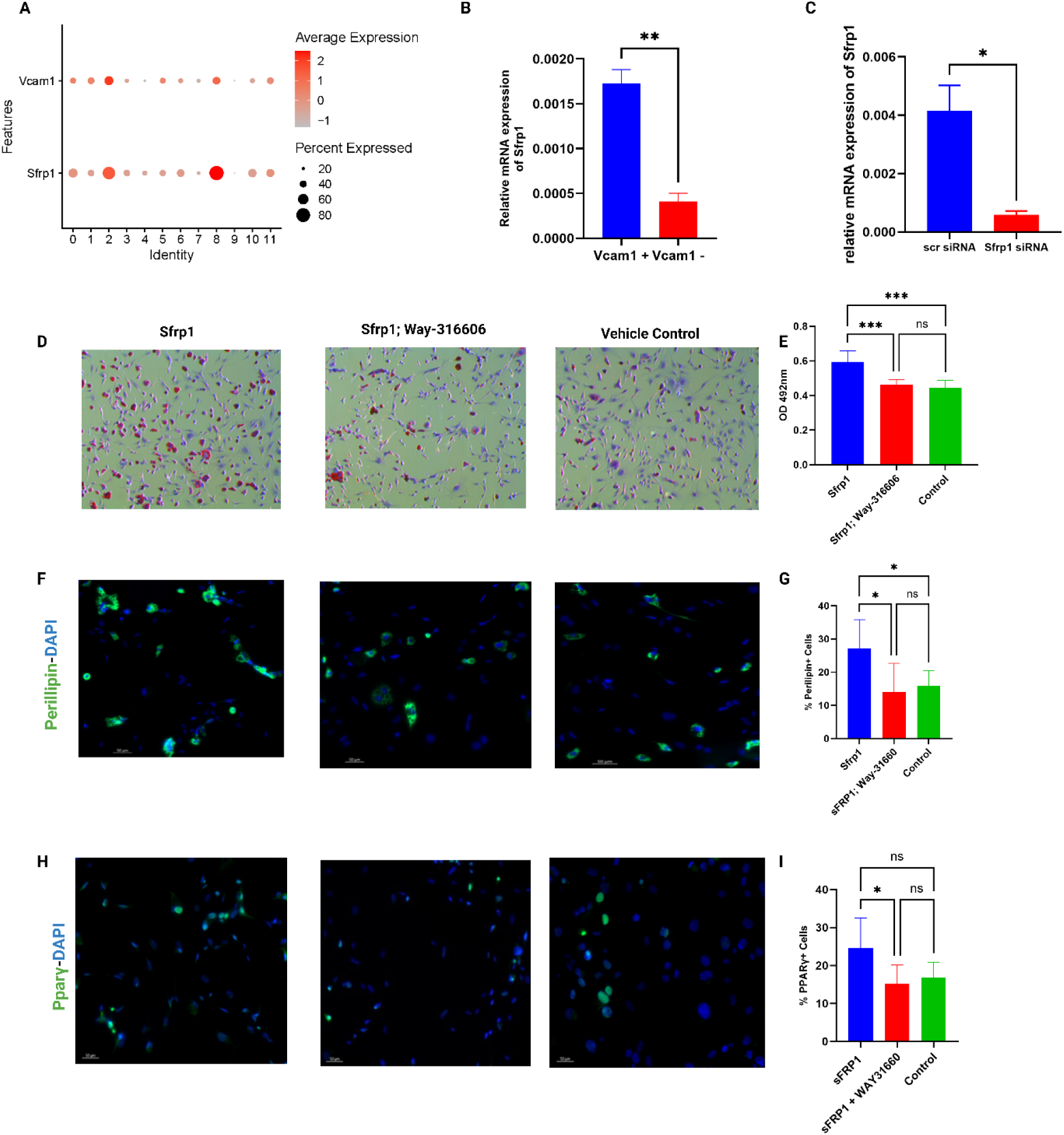
Sfrp1 regulates adipogenic differentiation in Vcam1+ FAPs (Related to Figure 4) A: Dot plot demonstrating Vcam1 and Sfrp1 expression in FAP subclusters B: Gene expression of Sfrp1 in Vcam1+ versus Vcam1-FAPs C: Gene expression of Sfrp1 in Sfrp1-silenced and control FAPs D: Representative images of ORO staining of Vcam1-FAPs treated with Sfrp1, way-316606, and a vehicle control E: Quantification of ORO staining of Vcam1-FAPs treated with Sfrp1, way-316606, and vehicle control. Data shown as mean ± SEM, ***p≤ 0.001, ns>0.05 F: Representative images of perilipin (green) and DAPI (blue) staining of Vcam1-FAPs treated with Sfrp1, way-316606, and a vehicle control G: Quantification of perilipin staining of Vcam1-FAPs treated with Sfrp1, way-316606, and vehicle control. Data shown as mean ± SEM, *p≤ 0.05, ns>0.05 H: Representative images of Pparγ (green) and DAPI (blue) staining of Vcam1-FAPs treated with Sfrp1, way-316606, and a vehicle control I: Quantification of Pparγ staining of Vcam1-FAPs treated with Sfrp1, way-316606, and vehicle control. Data shown as mean ± SEM, *p≤ 0.05, ns>0.05

**Supplementary Figure 4:**
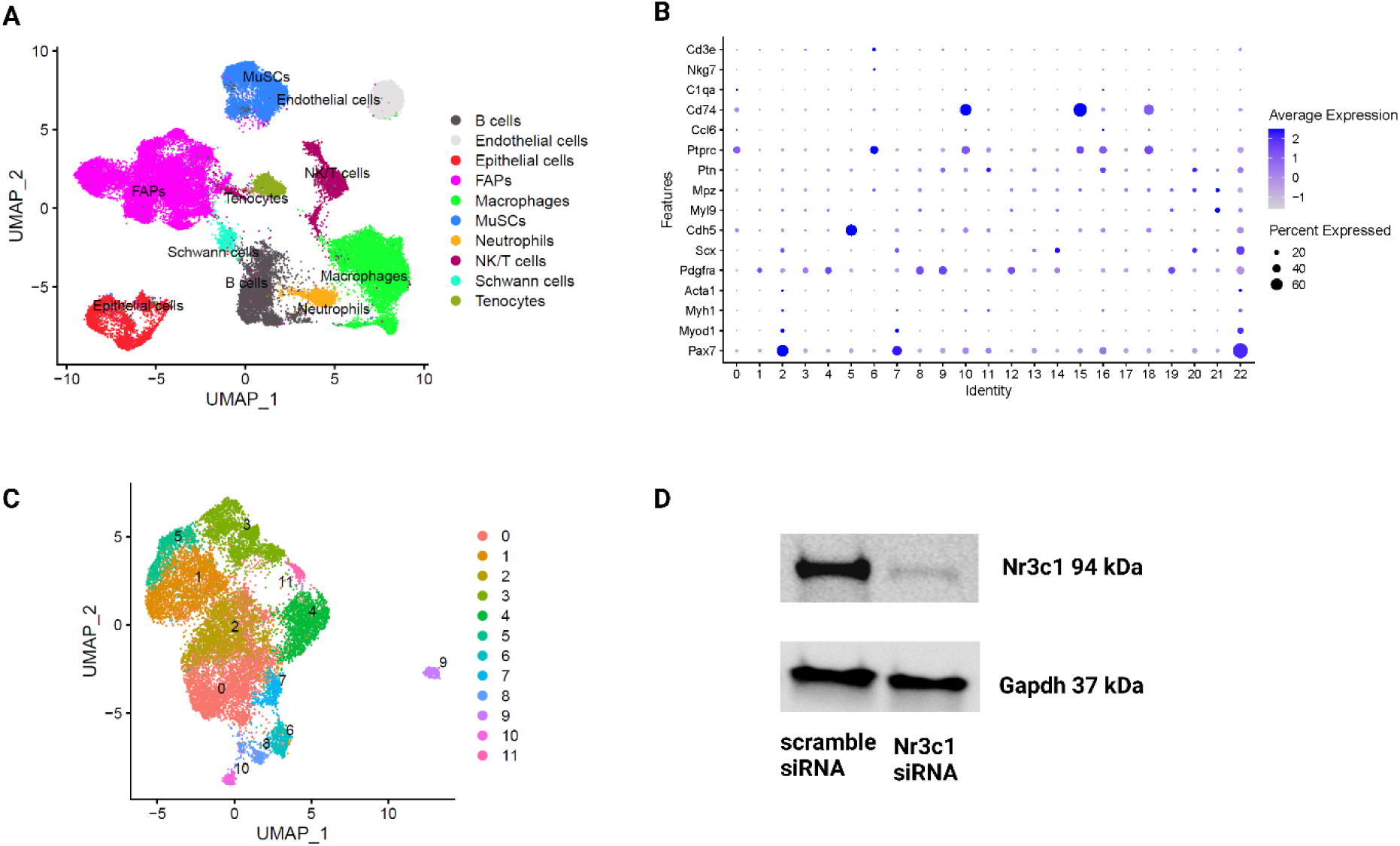
scATAC-seq reveals Nr3c1 as a candidate TF for adipogenic differentiation in Vcam1+ FAPs (Related to Figure 5) A: UMAP visualization of all cells, colored by cell type B: Dot plot demonstrating cell-type marker gene expression C: UMAP visualization of FAPs, colored by cluster D: Protein expression of Nr3c1 in Nr3c1-silenced and control FAPs

**Table S1:** Human PAD patient characteristics.

| Characteristics |  | All patients (N=8) |
| --- | --- | --- |
| **Demographics** |  |  |
| Mean patient age, years |  | 76 +/- 9.47 |
| Male sex (%) |  | 87.5 |
| Mean BMI |  | 30.44 +/- 5.67 |
| **Medical History (%)** |  |  |
| Diabetes |  | 87.5 |
| Hypertension |  | 100 |
| Hyperlipidemia |  | 100 |
| **Tobacco Use (%)** |  |  |
| Former smoker |  | 62.5 |
| Current smoker |  | 0 |
| **Medications (%)** |  |  |
| Statin |  | 100 |
| Aspirin |  | 62.5 |

